# Meru co-ordinates spindle orientation with cell polarity and cell cycle progression

**DOI:** 10.1101/2024.10.01.616065

**Authors:** Melissa M. McLellan, Birgit L. Aerne, Jennifer J. Banerjee Dhoul, Nicolas Tapon

## Abstract

Correct mitotic spindle alignment is essential for tissue architecture and plays an important role in cell fate specification through asymmetric cell division. Spindle tethering factors such as *Drosophila* Mud (NuMA in mammals) are recruited to the cell cortex and capture astral microtubules, pulling the spindle in the correct orientation. However, how spindle tethering complexes read the cell polarity axis and how spindle attachment is coupled to mitotic progression remains poorly understood. We explore these questions in *Drosophila* sensory organ precursors (SOPs), which divide asymmetrically to give rise to the epidermal mechanosensory bristles. We show that the scaffold protein Meru, which is recruited to the posterior cortex by the Frizzled/Dishevelled planar cell polarity complex, in turn recruits Mud, linking the spindle tethering and polarity machineries. Furthermore, Cyclin A/Cdk1 associates with Meru at the posterior cortex, promoting the formation of the Mud/Meru/Dsh complex via Meru and Dsh phosphorylation. Thus, Meru couples spindle orientation with cell polarity and provides a cell cycle-dependent cue for spindle tethering.

## Introduction

Accurate mitotic spindle orientation is key to development and adult homeostasis of multicellular organisms. In symmetrically dividing epithelial cells, the mitotic spindle generally aligns parallel to the tissue plane, allowing the daughter cells to insert themselves seamlessly into the epithelium, thereby maintaining tissue architecture and integrity (Bergstralh et al., 2017; Ragkousi & Gibson, 2014). Correct spindle orientation is also essential for the generation of cell diversity through the process of asymmetric cell division (ACD). During ACD, cell fate determinants (CFDs) are unequally inherited by the daughter cells, promoting progenitor self-renewal or differentiation (Sunchu & Cabernard, 2020a). CFD asymmetric segregation depends upon the polarisation of the cell cortex, which directs the spindle alignment and ensures unequal CFD inheritance. Thus, cell polarity and spindle orientation must be tightly linked, and aberrant alignment can result in developmental defects and neoplasia (Bergstralh et al., 2017; Ragkousi & Gibson, 2014; Sunchu & Cabernard, 2020b).

Mushroom body defective (Mud; Nuclear Mitotic Apparatus Protein/NuMA in mammals; LIN-5 in *C. elegans*) is a conserved spindle tethering factor that is recruited to the cell cortex by cell polarity proteins and exerts forces on astral microtubules by binding the Dynein/Dynactin motor complex (di Pietro et al., 2016). A core complex comprising the Mud binding partner Pins/LGN and Gαi has been implicated in Mud cortical recruitment (Bergstralh et al., 2017; Morin & Bellaïche, 2011; Siller & Doe, 2009). However, how Pins itself is recruited to the correct subcellular location and whether it is even required for Mud localisation is clearly context-dependent. For instance, *Drosophila* Mud localises to the cortex in a Pins-dependent manner in the ovarian follicular epithelium and via a Pins-independent mechanism in the wing and thoracic epithelia (Bergstralh et al., 2016; Bosveld et al., 2016; Nakajima et al., 2013, 2019; Neville et al., 2023). Thus, how the spatial alignment of the mitotic spindle adapts to the polarity of different symmetrically or asymmetrically dividing cell types remains unclear. Recent work has also proposed that, as well as being cortically recruited, the Mud/NuMA spindle tethering complex must be activated as the cells enter mitosis (Neville et al., 2023). However, our understanding of how the spindle tethering machinery is coupled to cell cycle progression remains limited. Here, we use the asymmetric division of *Drosophila* sensory organ precursor cells (SOPs) as a model to study these questions.

SOPs (also known as pI), which give rise to the adult sensory bristles of the fly epidermis, are a well-studied example of ACD (Hartenstein & Posakony, 1989; Schweisguth, 2015). The best-studied SOPs are located on the dorsal thorax (notum) and produce the mechanosensory microchaetes that cover this tissue (Hartenstein & Posakony, 1989; Schweisguth, 2015). SOPs undergo consecutive rounds of asymmetric division to produce five distinct cell types: neuron, sheath, shaft, socket and a glial cell that undergoes apoptosis (Hartenstein & Posakony, 1989). The SOP lineage cells then recruit a neighbouring epidermal cell (the F-Cell) and together these assemble into a functional sensory hair(Mangione et al., 2023). In the notum, SOPs are specified at ∼12 h APF (hours After Puparium Formation) from the epithelial sheet through the highly conserved Notch pathway (Corson et al., 2017; Gómez-Skarmeta et al., 2003; Simpson, 2007).

Until the SOPs are specified, they share the same polarity as the surrounding epithelial cells, characterised by two polarity axes: (1) planar cell polarity (PCP), whereby the cells align in the tissue plane and (2) and apical-basal (A-B) polarity which defines the cellular apical and basal domains separated by the adherens junctions (AJs) (Buckley & St Johnston, 2022; Goodrich & Strutt, 2011). The core PCP pathway is established by three transmembrane proteins that form opposing domains at the AJs through mutual antagonism (Goodrich & Strutt, 2011; Yang & Mlodzik, 2015). In the notum, heterodimers of the seven-pass transmembrane protein Flamingo (Fmi, also known as Starry night/Stan; CELSR1/2/3 in vertebrates) with the four-pass transmembrane protein Van Gogh (Vang, also known as Strabismus; Vang1/2 in vertebrates) are located on the anterior side of each cell, while heterodimers of Fmi with the seven-pass protein Frizzled (Fz; Fz1 in vertebrates) are on the posterior side (Bellaïche, Gho, et al., 2001; Schweisguth, 2015; Ségalen et al., 2010). Vang then recruits its downstream effector Prickle (Pk) (Bastock et al., 2003; Jenny et al., 2003; Taylor et al., 1998), while Fz recruits Dishevelled (Dsh; DVL1-3 in vertebrates) and Diego (Dgo) (Axelrod et al., 1998; Feiguin et al., 2001; Jenny et al., 2005). In A-B polarity, the apical domain is defined by the Par complex, comprised of Bazooka (Baz; Par3 in vertebrates), atypical Protein Kinase C (aPKC; PKCι/PKCζ in vertebrates) and Par6 which regulates the placement of the adherens junctions (Bilder et al., 2000; Buckley & St Johnston, 2022; Tepass, 2012). The basolateral domain is defined by the septate junction components Scribble (Scrib; SCRIB in vertebrates), Discs large (Dlg; Dlg1-5 in vertebrates) and Lethal giant larvae (Lgl; LLGL1/2 in vertebrates) (Albertson et al., 2004; Bilder et al., 2000; Woods & Bryant, 1991).

Upon SOP specification, the proneural transcription factors of the Achaete-Scute complex turn on the expression of the N-terminal RASSF (Ras association domain family) protein Meru (Banerjee et al., 2017; Buffin & Gho, 2010; Reeves & Posakony, 2005). Meru is recruited to the posterior cortex by Dsh, and in turn recruits Baz, leading to its planar asymmetry specifically in the SOP (Banerjee et al., 2017). Baz then polarises the rest of the Par complex to the posterior cortex, leading to exclusion of the CFDs Numb and Neuralized (both Notch pathway modulators) from the posterior pole prior to ACD (Bellaïche, Gho, et al., 2001; Bellaïche, Radovic, et al., 2001; Besson et al., 2015; Le Borgne & Schweisguth, 2003; Roegiers et al., 2001a, 2001b; Smith et al., 2007; Wirtz-Peitz et al., 2008). Thus, Meru provides the initial planar bias that triggers polarisation of the Par complex and the CFDs along the antero-posterior axis (Banerjee et al., 2017). To ensure accurate segregation of CFDs, the mitotic spindle must also align along the antero-posterior axis. This is achieved by redundant spindle tethering complexes on the anterior and posterior poles (Gomes et al., 2009; Schweisguth, 2015) (Figure 1A). On the anterior side, Gαi/Dlg/Pins recruit Mud basally, while on the posterior side, Mud is recruited in a Fz/Dsh-dependent manner to the apical cortex (Bellaïche, Gho, et al., 2001; Bellaïche, Radovic, et al., 2001; David et al., 2005; Johnston et al., 2013; Schaefer et al., 2000; Ségalen et al., 2010). This allows the capture of one centrosome each by the anterior and posterior poles, but also imparts a characteristic A-B tilt to the spindle (David et al., 2005; Schweisguth, 2015; Ségalen et al., 2010) (Figure 1A). Thus, SOP division is a powerful system to study how spindle orientation adapts to cellular context, since two distinct Mud localisation mechanisms co-exist in this cell type.

**Figure 1.**
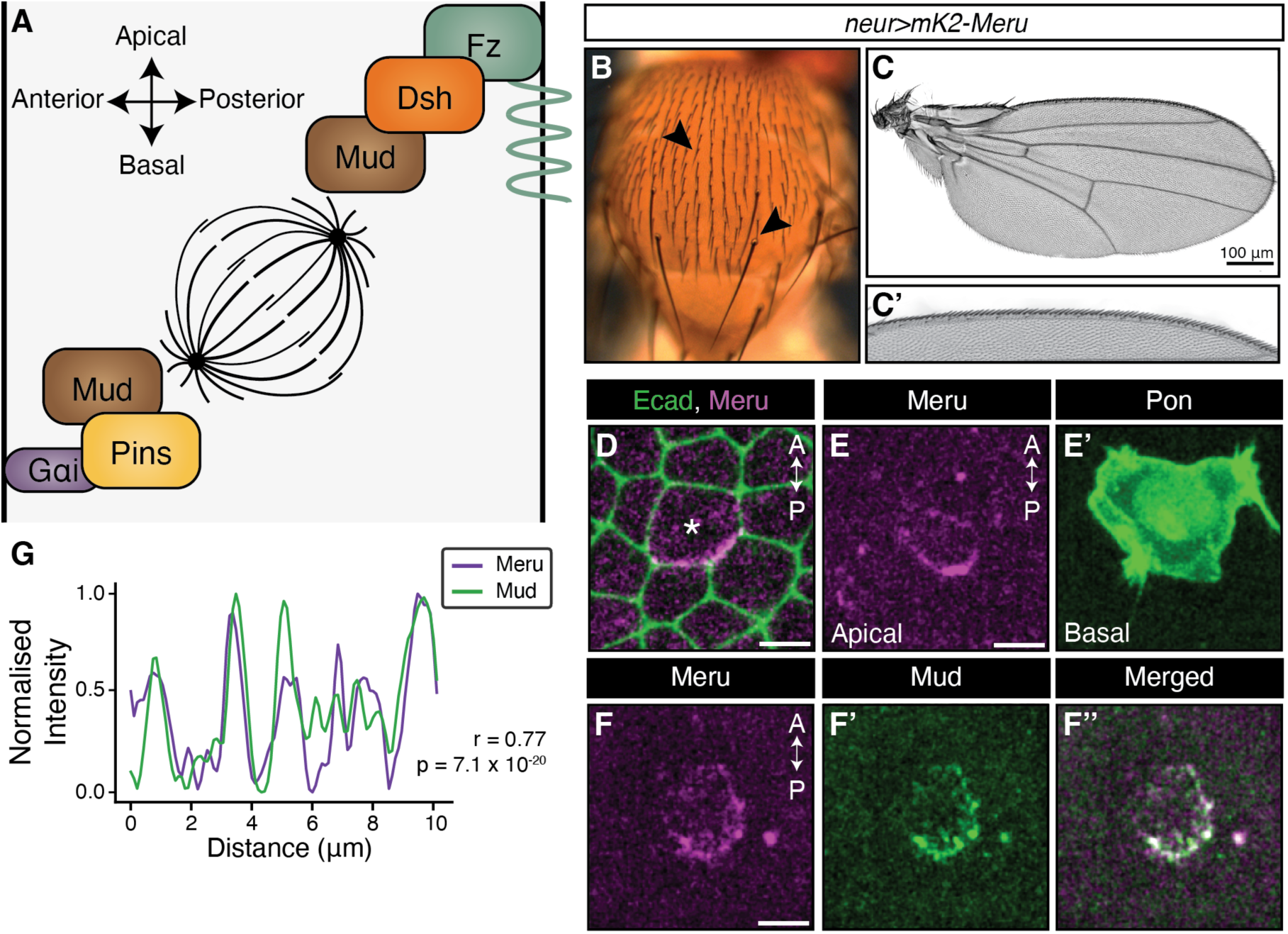
Meru localises to the apical posterior domain of the SOP. (**A**) The current model of SOP spindle alignment where Mud is recruited to the anterior cortex by Pins/G*α*i and to the posterior cortex through Fz/Dsh. (**b, c**) Brightfield images of the adult notum (**B**), wing (**C**), and anterior wing margin (**C’**) in animals expressing *UAS-mK2-meru* under the *neurG4* driver. No visible defects (arrows indicating micro/macrochaetes) in the notum and wing sensory organs were observed. (**D, E, F**) Maximum intensity projections of pupal nota at 16 h APF of *neurG4>UAS-mK2-meru* (magenta, **D, E** and **F**), *Ecad-GFP*, *Pon-GFP* and *mud-GFP* (green; panels **D**, **E’** and **F’**, respectively) at 18 mins prior to first indication of cytokinesis. SOP marked by star. (**G**) Single brightest slice in panels **F** and **F’** plotted as line graph by measuring the gray value across the posterior domain as defined by Meru localisation normalised to highest value in each channel. High Pearson’s coefficient (*r*) indicates positive correlation. p-value calculated using a two-tailed test. Scale bars = 5 µm.

An important open question remains how Mud is recruited to the posterior cortex as the SOPs enter mitosis. Two potential mechanisms have been proposed: (1) Mud binds directly to Dsh (Ségalen et al., 2010) and (2) the mitotic cyclin, Cyclin A (CycA), is recruited to the posterior cortex in a Dsh-dependent manner and in turn recruits Mud (Darnat et al., 2022). Here, we show that, in parallel to its role in posterior Baz recruitment, Meru acts as a bridging factor between Dsh and Mud. Examination of the Meru sequence revealed a CycA docking site that is required for Meru cortical localisation, Mud localisation and spindle alignment. We also identified multiple Cdk1 phosphorylation consensus sites that are essential for the Meru/Mud interaction. Meru therefore links PCP with spindle tethering, and its ability to recruit Mud is coupled to the cell cycle through CycA to ensure timely alignment of the spindle during mitosis.

## Results

### Meru localises to the apical-posterior pole prior to and during SOP mitosis

To address the potential role of Meru in SOP spindle alignment, we generated a *UAS-mKate2(mK2)-meru* transgenic line to track its localisation during SOP divisions relative to known polarity and spindle factors. We had previously shown that endogenously tagged GFP-Meru localises to the posterior cortex at interphase, where it remains enriched throughout mitosis and co-localises with Dsh (Banerjee et al., 2017). No bristle defects were observed in *neurG4; mK2-meru* animals, suggesting overexpression does not lead to strong SOP division defects (Figure 1B-C’). As expected (Banerjee et al., 2017), mK2-Meru driven in SOPs by the *neur^P72^-GAL4* driver (*neurG4*) (Bellaïche, Gho, et al., 2001) localised to the posterior apical cortex at the level of E-cadherin (Ecad) (Figure 1D and EV1A). Upon mitotic rounding, Partner of Numb (Pon) is rapidly relocalised to the basolateral anterior cortex during prophase, followed by its unequal inheritance into the anterior pIIb daughter cells (Bellaïche, Gho, et al., 2001; Roegiers et al., 2001b). Co-expression of mK2-Meru and Pon showed that mK2-Meru was present at the posterior domain before the asymmetric localisation of Pon was detectable, indicating SOP polarisation occurs prior to CFD segregation (Figure 1E-E’). Thus, Meru is localised at the posterior apical cortex during most of SOP mitosis, consistent with a possible role in spindle positioning.

### Meru is required for the cortical recruitment of Mud via Dsh

Dsh and Mud have been shown to co-localise at the posterior-apical cortex during SOP mitosis (Ségalen et al., 2010). We reported that Meru and Dsh associate in cell culture, co-localise in fixed tissue, and that Dsh is required for Meru posterior localisation (Banerjee et al., 2017). We therefore sought to test whether there was an interaction between Meru and Mud by tracking the localisation of mK2-Meru and a GFP-tagged Mud construct expressed under the *mud* promoter (GFP-Mud) with live-imaging (Ségalen et al., 2010). Mud and Meru localisations were highly correlated (as indicated by a Pearson’s coefficient *r* near +1), particularly at the onset of mitosis (Figure 1F-G). Despite a considerable drop in Mud intensity levels, the two proteins remained positively correlated even after metaphase (Fig EV1B, B’ and C, C’).

The strong Meru/Mud colocalisation is consistent with a physical interaction between these proteins. To test this possibility, we performed co-immunoprecipitations (co-IPs) from S2 cell lysates expressing full-length Meru together with Mud fragments spanning much of the protein (Ségalen et al., 2010) (Figure 2A). We found that, despite its low expression, the most C-terminal fragment Mud(1825-2457), which contains a small portion of the coiled-coil region but is largely disordered, strongly associated with Meru (Figure 2B). Interestingly, this disordered region also includes a short domain that mediates Pins binding, known as the Pins Binding Domain (PBD), and is required to recruit Mud cortically in SOPs and neuroblasts (Ségalen et al., 2010; Siller et al., 2006). To test if the PBD was also essential for the Mud/Meru interaction, the Mud(1452-1824) non-binding fragment was extended N-terminally to include the PBD (Mud(1452-1961)). Addition of the PBD was sufficient to confer Meru binding (Figure 2B), suggesting that Mud interacts with Meru via the C-terminal PBD.

**Figure 2.**
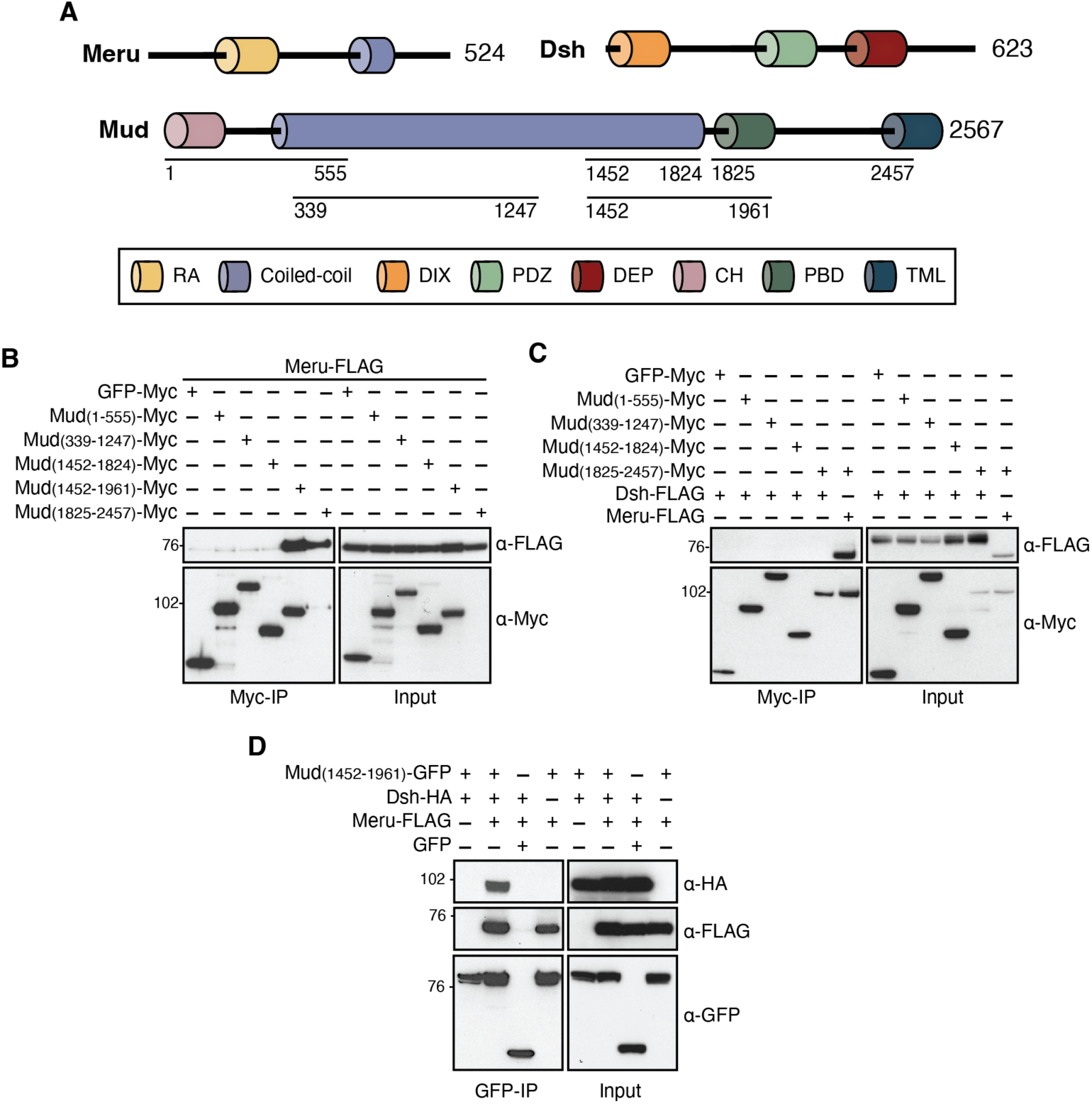
Meru is required for Dsh/Mud complex formation. **(A**) Schematic of constructs that span Meru, Dsh and Mud used in co-IPs to characterise Meru binding in S2 cells. (**B, C, D**) Western blots of co-IP experiment using S2 cell lysates from transfected S2 cells, immunoprecipitated using and probed with the indicated antibodies. (**B**) Full-length Meru immunoprecipitates with Mud fragments containing the PBD. (**C**) C-terminal fragment of Mud does not co-IP with full-length Dsh. The Meru/Mud interaction is used as a positive control (right-most lane). (**D**) Meru promotes Dsh/Mud complex formation.

Previous reports showed an interaction between Mud and Dsh in HEK293T cells (Ségalen et al., 2010). We were unable to detect such an interaction in S2 cells between full-length Dsh and any of the Mud fragments (Figure 2C). However, Meru is not endogenously expressed in S2 cells, whereas the orthologs of Meru (RASSF9/10) are expressed in HEK293T cells (Hauri et al., 2013). Together with our finding that Meru interacts with both Dsh and Mud in cell culture, this prompted us to test whether Meru could bridge the previously reported Dsh/Mud interaction (Ségalen et al., 2010). Indeed, co-expression of Meru elicited a robust association between Dsh and Mud(1452-1961) (Figure 2D). Thus, Meru co-localises with Mud and Dsh at the posterior cortex during SOP cell divisions and mediates the assembly of a Mud/Meru/Dsh complex in cell culture.

### Meru is required for cortical Mud localisation and spindle orientation *in vivo*

In wild-type SOPs and epithelial cells, Mud localises to three regions: at the cortex (with polarity proteins), at the centrosomes, and at tricellular junctions (Bosveld et al., 2016; Izumi et al., 2006; Ségalen et al., 2010). Specifically, Mud localisation to the posterior cortex in SOPs is dependent on Dsh (Ségalen et al., 2010). As Meru is required for the Dsh/Mud interaction in cell culture, we tested whether posterior cortical Mud recruitment is dependent on Meru. We quantified Mud-GFP at both the anterior and posterior cortex in *meru^1^* mutants compared to wild-type flies (Figure 3A-C). We noted that, in wild-type SOPs, Mud crescent intensity is consistently higher on the anterior side than the posterior (Figure 3C). We observed a significant decrease in Mud intensity at the posterior cortex of *meru^1^* mutant SOPs relative to the wild type (Figure 3C). However, anterior cortical Mud levels were also decreased, potentially indicating a role for Meru in Mud stability (Figure 3C). Thus, Meru is required for Mud cortical recruitment, consistent with a role in spindle alignment.

**Figure 3.**
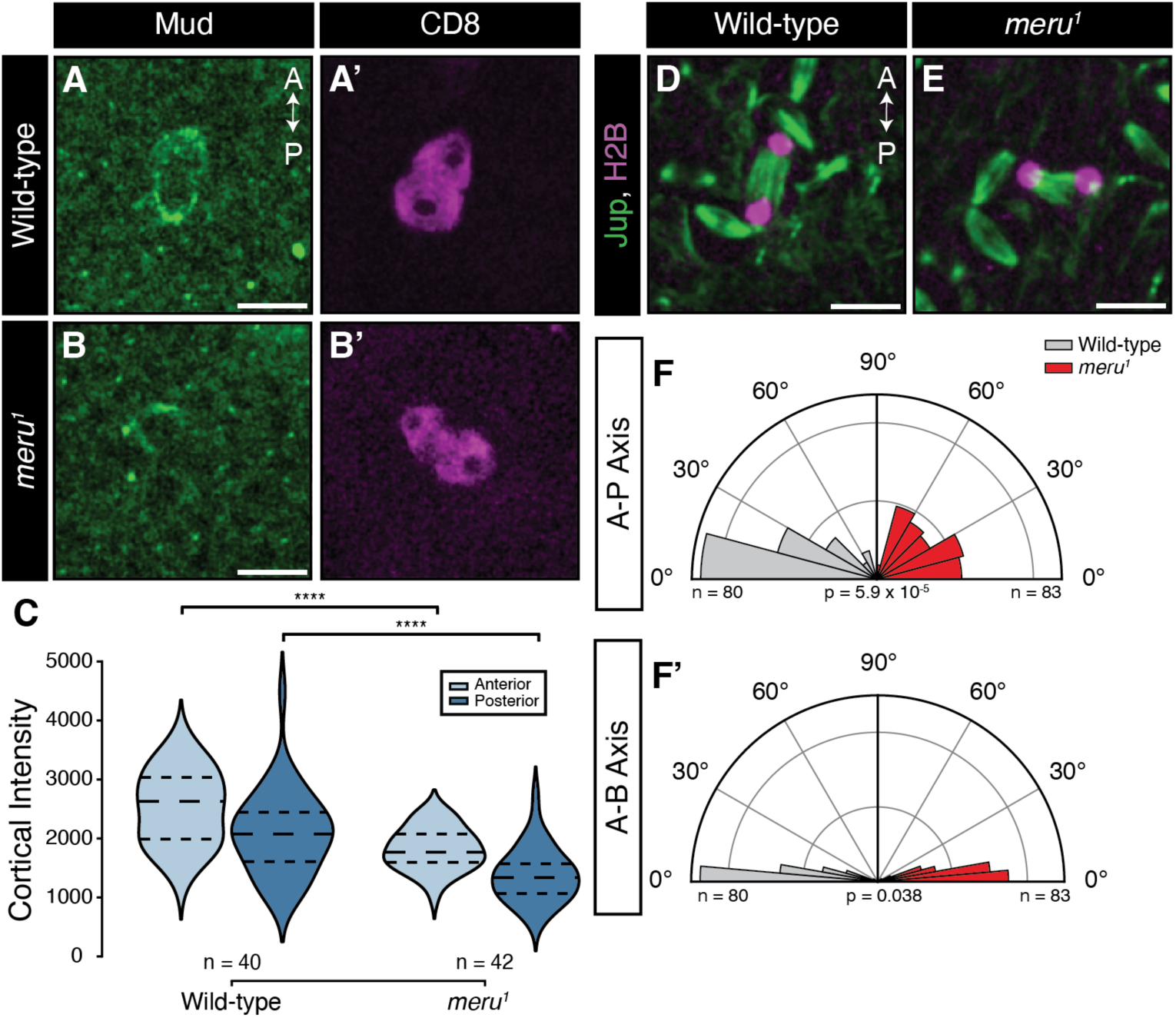
Meru loss leads to Mud mislocalisation and spindle alignment defects. (**A, B**) Pupal notal confocal live-imaging of *Mud-GFP* (green) and *UAS-cd8-mRFP* driven by *neurG4* (magenta) in a *meru* wild-type (**A**) and *meru^1^* background (**B**) at 16 h APF at the first indication of cytokinesis. (**C**) Graph showing the intensity of cortical Mud at the anterior and posterior crescent of each genotype. Both the anterior and posterior cortical Mud levels were significantly lower in the *meru^1^* flies relative to the wild-type (p = 1.4 x 10^-6^ and p = 1.4 x 10^-6^, respectively). **** = p < 0.0001 using a Mann-Whitney U test. (**D, E**) Confocal live-imaging of *neur-H2B-RFP* (magenta) and *Jupiter-GFP* (green) in a *meru* wild-type (**D**) and *meru^1^* background at approximate cytokinesis (**E**). (**F**) Graphs showing the spindle orientation of each genotype relative to the dorsal midline (**F**) and epithelial plane (**F’**). p-values indicated using a Kolmogorov-Smirnov test. Scale bars = 10 µm. Number of SOPs imaged indicated on the panels from 3 animals in panel (**C**) or 6 wild-type or 4 *meru^1^* animals, respectively, in panels (**F-F’**).

We previously showed that *meru* mutant SOPs display spindle orientation defects (Banerjee et al., 2017). However, these measurements were performed in the presence of overexpressed *pon*, which is known to induce changes in SOP polarity (Perdigoto et al., 2008). We therefore directly measured spindle alignment by labelling the spindle using a GFP-tagged version of the microtubule-binding protein Jupiter and quantifying the deviation from the A-P axis in wild-type and *meru^1^* mutant flies (Figure 3D-F’, Movie EV1 and EV2). As the mitotic spindle remains dynamic until metaphase, when it settles for its final division angle (Bellaïche, Gho, et al., 2001; Bergstralh et al., 2016; Ségalen et al., 2010), our measurements were carried out post-metaphase. Consistent with previous work (Ségalen et al., 2010), 71% of wild-type cells divided within 30° of the A-P axis (Figure 3D and F), and the spindle alignment in the A-B axis averaged at 6.8°, displaying the characteristic z-tilt of SOP mitotic spindles (David et al., 2005; Ségalen et al., 2010) (Figure 3F’). Strikingly, *meru^1^* flies had a nearly random spindle alignment in the A-P axis, with only 45% of cells dividing within 30° of the midline (Figure 3E and F). The average A-B angle showed a slight increase to 9.0°, with a significantly wider distribution – nearly 20% of cells dividing over 20°, compared to 2*%* in controls (Figure 3F’). Consistent with its role in Mud positioning, Meru is therefore required for A-P and planar spindle alignment.

### Meru associates with CycA

CycA has recently been reported to be enriched at the apical posterior cortex at the end of G2 and early prophase in SOPs (Darnat et al., 2022). We therefore wished to investigate whether CycA could provide a link between cell cycle progression and spindle orientation by functioning in concert with Meru. We first tracked Meru/CycA colocalisation *in vivo*. In late G2 phase, when both proteins have been reported to be present at the posterior cortex during SOP mitosis (Banerjee et al., 2017; Darnat et al., 2022), we observed that the localisations of mK2-Meru and endogenously tagged CycA were highly correlated (Figure 4A-B – Pearson’s coefficient = 0.71). The colocalisation persisted until metaphase (Fig 4A), when CycA enters the nucleus, which is followed by its degradation (Lehner & O’Farrell, 1989). We then tested a potential Meru/CycA association by co-IP in S2 cells. Interestingly, CycA robustly immunoprecipitated with Meru, but not Dsh or Mud (Figure 4C). To determine whether the Meru/CycA interaction is specific, we examined binding to other Cyclins associated with cell cycle progression (A, B, E and D). We observed an interaction with CycB, which is expressed during early M-phase (Lehner & O’Farrell, 1990). However, this CycB association was far weaker than the interaction with CycA (Figure 4D). Thus, Meru binds to CycA, with which it co-localises at the posterior cortex in SOPs.

**Figure 4.**
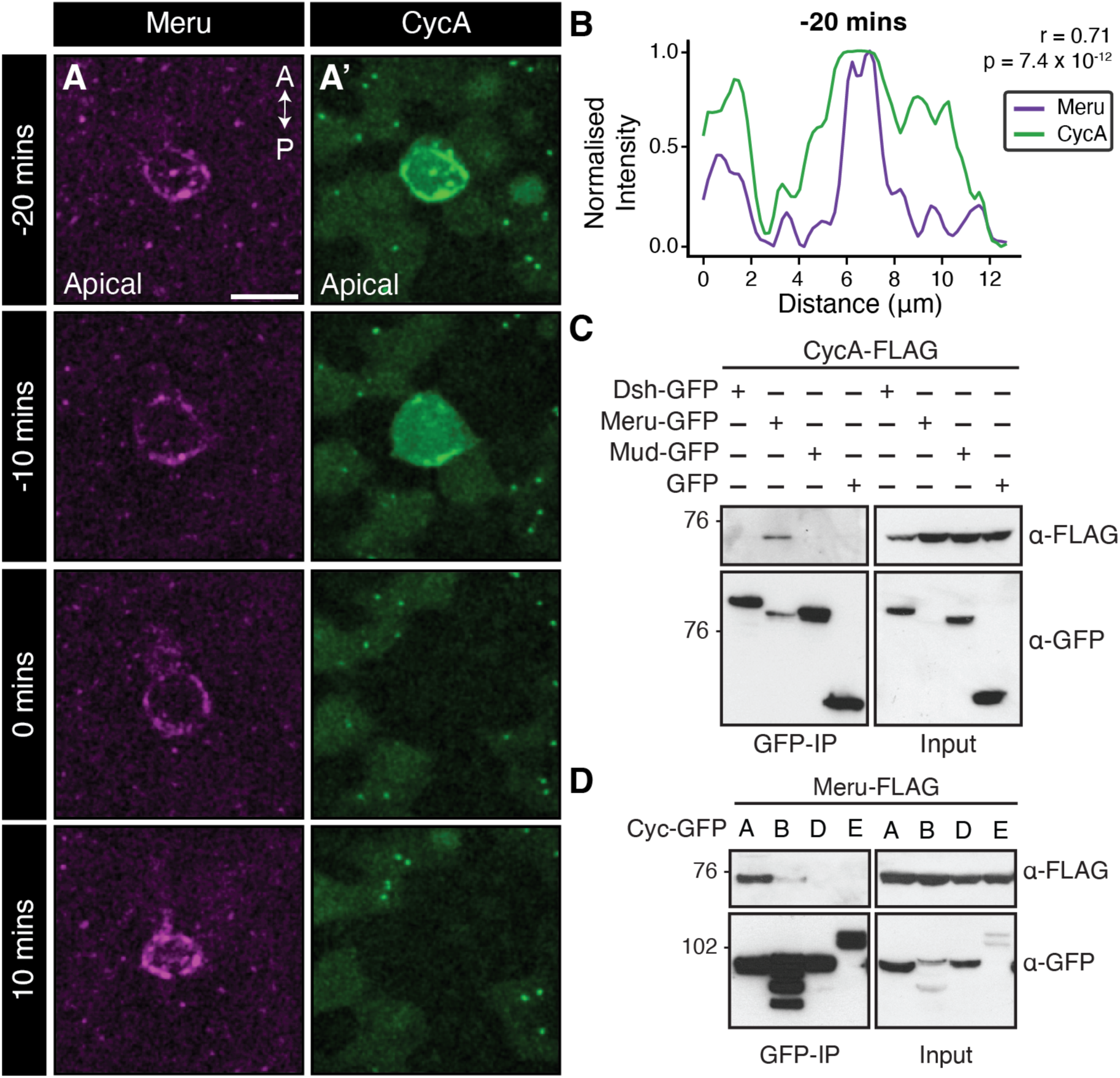
Meru and CycA interact at the posterior cortex of SOPs. **(A**) Confocal live-imaging of pupal nota expressing *neurG4>UAS-mK2-meru* (magenta) and *CycA-GFP* (green) of an SOP during mitosis at 16 h APF and 0 mins indicating the first frame in cytokinesis. (**B**) Single brightest slice in panels **A** and **A’** at −20 mins plotted as line graph by measuring the gray value across the posterior domain as defined by Meru localisation. High Pearson’s coefficient (*r*) indicates positive correlation. p-value calculated using a two-tailed test. (**C, D**) Western blots of co-IP experiment using cell lysates from transfected S2 cells, immunoprecipitated using *α*-GFP beads and probed using *α*-FLAG and *α*-GFP antibodies. (**C**) Meru, not Dsh or C-terminal Mud_(1452-1961)_, immunoprecipitates with CycA. (**D**) Meru associates with CycA and weakly with CycB, but not CycD or CycE. Scale bar = 10 µm.

### CycA docking regulates Meru localisation and association with Mud and Dsh

CycA binding is highly regulated through short linear motifs (or SLiMs), the most well-known of which is the RxL motif. This consensus sequence consists of amino acids R/K-x-L-φ or R/K-x-L-x–φ (φ = hydrophobic amino acid) (Tatum & Endicott, 2020) (Fig EV2). When we investigated the Meru sequence, we found six possible RxL motifs, two of which (RxL 258 and 259) were very good matches and located immediately adjacent to each other (Figure 5A and Fig EV2). We generated mutations at all six RxL motifs, substituting the R/K and L to A. We initially found that Meru/CycA binding was only affected in the RxL 258/9 double mutant (Fig EV3). Subsequent mutation of either the 258 or 259 site alone showed that this was sufficient to disrupt the Meru/CycA interaction (Figure 5B). In all subsequent experiments, we used the double 258/9 mutants (*meru^RxL^*).

**Figure 5.**
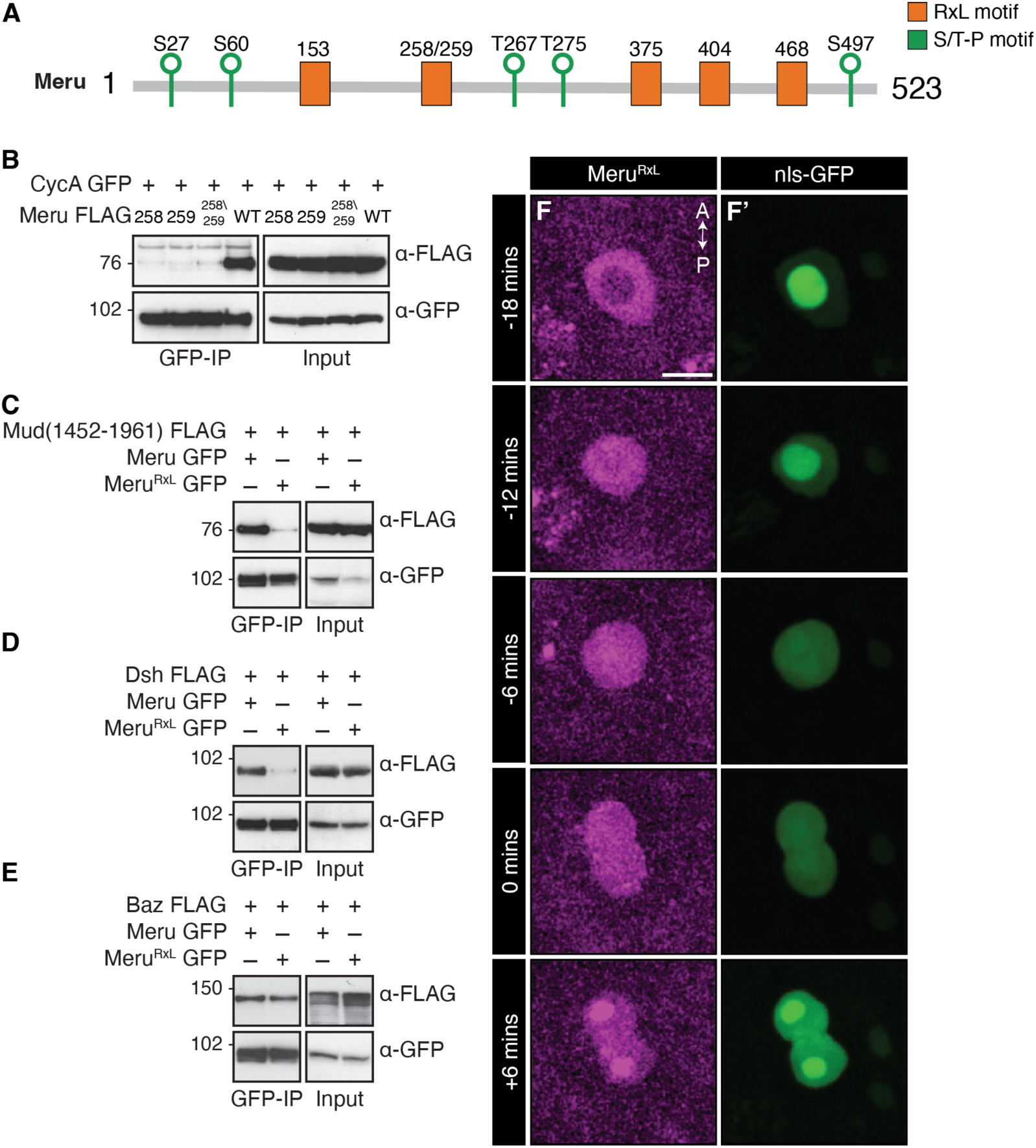
CycA docking is required for Meru localisation *in vivo*. **(A)** Schematic of RxL motifs and Cdk1 S/T-P phosphorylation sites present in the Meru sequence. **(B)** Meru mutated at RxL motifs 258, 259 and 258/9 no longer immunoprecipitates with CycA. (**B, C, D, E**) Western blots of co-IP experiment using cell lysates from transfected S2 cells, immunoprecipitated and probed using the indicated antibodies. Mutation of Meru RxL 258 and 259 sites disrupt the interaction between Meru and CycA (**B**), Mud (**C**) and Dsh (**D**), but not Baz (**E**). (**F**) Confocal live-imaging of pupal nota at 16 h APF and 0 mins marking first frame in cytokinesis shows expression of *UAS-mK2-meru* (magenta) and *UAS-nls-GFP* (green) driven by *neurG4* during SOP mitosis. Scale bar = 10 µm.

To determine if CycA binding modulates Meru function, we tested if the RxL 258/9 motif was required for association with other Meru partners. We observed that the Meru RxL motif is necessary for Dsh and Mud binding (Figure 5C and D), but dispensable for Baz (Figure 5E). To distinguish between the RxL mutation directly interfering with Mud binding versus Meru/CycA binding being required for Meru to associate with Mud, we depleted CycA by RNAi (Fig EV3B). CycA expression was required for the Meru/Mud association, consistent with the RxL mutation interfering with Meru/Mud binding indirectly by compromising CycA docking to Meru.

We then generated *UAS-meru^RxL^* transgenic flies to test the role of Meru/CycA binding *in vivo*. Strikingly, mK2-Meru^RxL^ failed to be recruited to the cortex, in agreement with the RxL mutation abolishing Dsh binding (Figure 5F-F’, compare with Fig 1 and Fig EV1). We also noted that, after nuclear envelope reassembly, some mK2-Meru^RxL^ accumulated in the nucleus. Co-expression with Ecad and Dlg showed that mK2-Meru^RxL^ was distributed in both the apical and basal cytoplasm (Fig EV3C-D’). In summary, CycA docking to Meru via the RxL motif is required for Dsh and Mud binding in cell culture, as well as Meru cortical localisation *in vivo*.

### Meru Cdk1 phosphorylation sites modulate binding to Mud and Dsh

As CycA’s best characterised function is Cdk1 regulation, we investigated whether Meru phosphorylation by Cdk1 could affect Meru/Mud association. Upon CycA/Cdk1 co-expression in S2 cells, we observed a marked increase in Meru/Mud binding compared to basal levels (Fig EV4A). The minimal Cdk1 phosphorylation consensus site is S/T-P, while the optimal consensus is S/T-P-x-K/R (Enserink & Kolodner, 2010). We identified five potential phosphorylation sites within the Meru sequence (Figure 5A). Although several phosphosite mutants affected the Meru/Mud association, no single mutation was sufficient to abolish the interaction (Fig EV4B). However, when all five sites were mutated (Meru^5S/T-A^) Meru/Mud binding was eliminated (Figure 6A). In contrast, Dsh and Baz could still interact with the Meru^5S/T-A^ mutant (Figure 6A). Examining the Dsh sequence, we found a single Cdk1 phosphosite at amino acid 22 within the DIX (DIshevelled and aXin) domain. Interestingly, mutating this site (Dsh^T22A^) had no impact on binding to wild-type Meru (Figure 6B). However, when the Dsh^T22A^ mutant was combined with the Meru^5S/T-A^ mutant, the association was almost completely lost. These data suggest that Meru phosphorylation by Cdk1 is required for Meru/Mud binding, while Meru/Dsh assembly is dependent on phosphorylation of both proteins.

**Figure 6.**
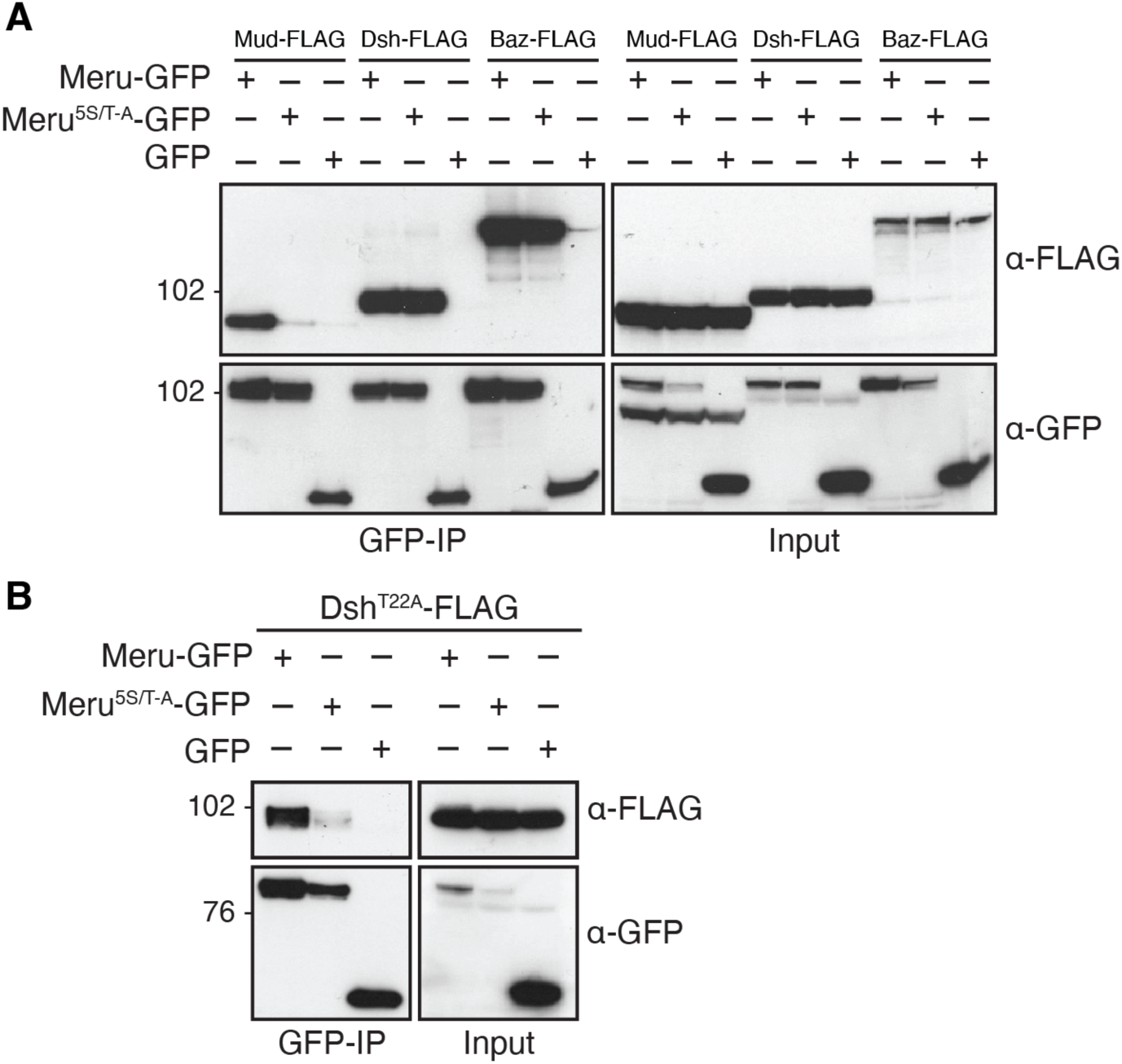
Phosphorylation by Cdk1 is required for Meru binding to Mud and phospho-Dsh. **(A, B)** Western blots of co-IP experiment using cell lysates from transfected S2 cells, immunoprecipitated and probed using the indicated antibodies. Meru mutated to Alanine at five S/T-P phosphorylation sites disrupts co-immunoprecipitates with C-terminal Mud_(1452-1961)_, but not Baz or Dsh (**A**), unless the Dsh T22 phosphorylation site was also mutated (**B**).

### CycA docking is required for Meru-dependent spindle orientation

We sought to address if the role of Meru in SOP spindle alignment was regulated by CycA. We first tested the effect of blocking Meru/CycA association on the formation of the ternary Dsh/Meru/Mud complex in S2 cells. In contrast to wild-type Meru, the Meru^RxL^ mutant was unable to bridge the Dsh/Mud interaction (Figure 7A), suggesting that CycA binding to Meru is required for ternary complex formation. Second, we tested if loss of CycA binding impacted Meru’s role in spindle orientation *in vivo*. We performed a rescue experiment by expressing *UAS-mK2-meru* constructs in a *meru^1^* mutant background and quantified cortical Mud localisation as in wild-type SOPs above (Figure 3D-F’). In the rescued conditions, both the anterior and posterior cortices had nearly identical levels of Mud when mK2-Meru^WT^ was expressed (Figure 7B, B’ and D). It is interesting to note that, in the absence of Meru overexpression, Mud is more enriched at the anterior than the posterior cortex (Fig 3C). The equal cortical distribution of Mud upon mK2-Meru^WT^ overexpression is therefore consistent with Meru promoting Mud posterior recruitment. In contrast, mK2-Meru^RxL^-expressing SOPs had reduced Mud levels at the posterior cortex (Figure 7C, C’ and D), suggesting that CycA binding of Meru is necessary for posterior Mud localisation.

**Figure 7.**
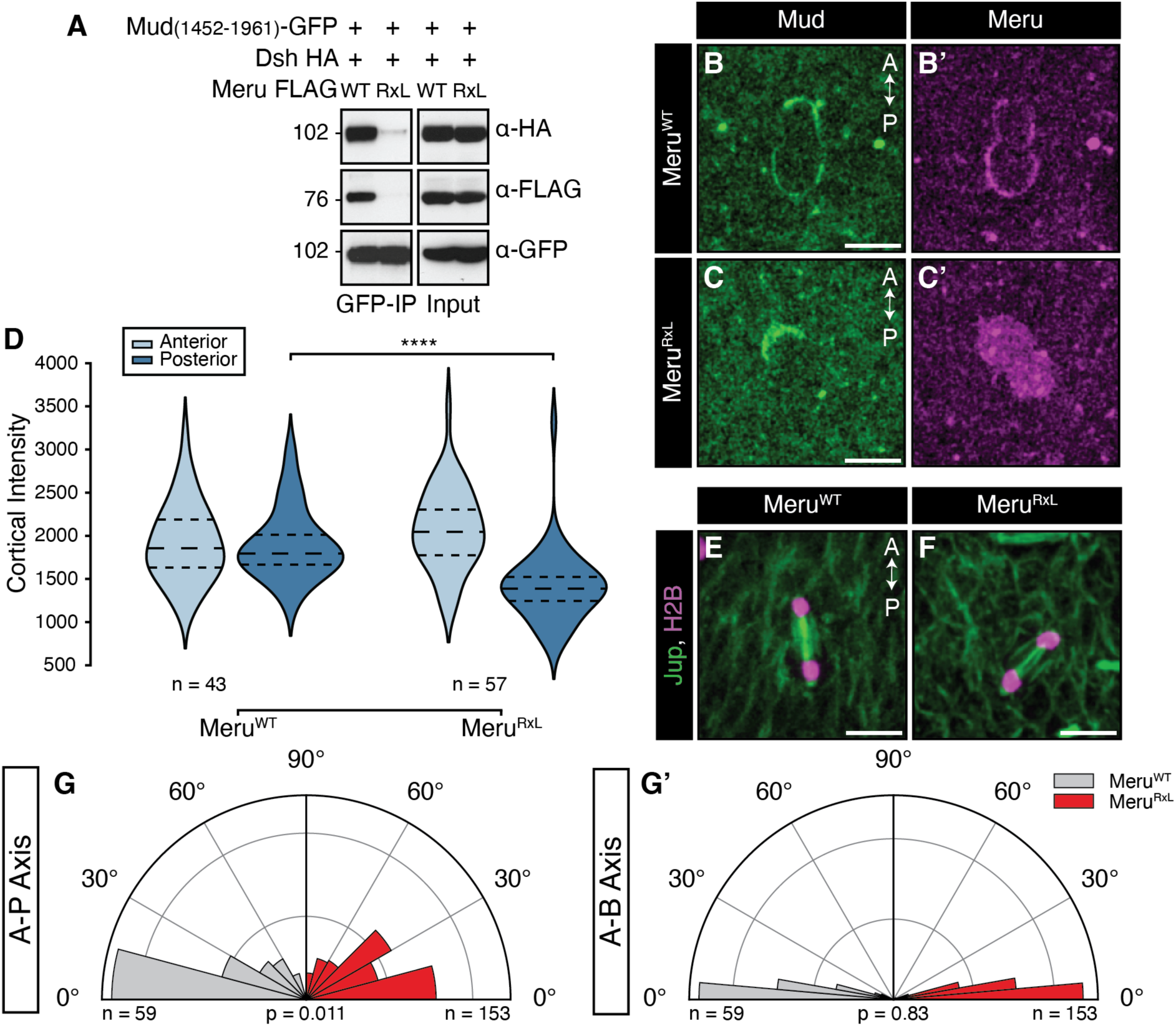
Mutation of the Meru RxL motif results in decreased posterior Mud and abnormal spindle alignment. **(A)** Western blot of co-IP experiment using cell lysates from transfected S2 cells, immunoprecipitated and probed using the indicated antibodies. Dsh does not immunoprecipitate with C-terminal Mud in the presence of Meru^RxL^. (**B, C**) Confocal live-imaging of pupal nota at 16 h APF expressing *Mud-GFP* (green) and *neurG4>UAS-mK2-meru^WT^* (**B**) or *neurG4>UAS-mK2-meru^RxL^* (**C**) in a *meru^1^* background. (**D**) Graph showing the intensity of cortical Mud at the anterior and posterior crescent of each genotype. Posterior *meru^RxL^* is significantly lower than *meru^WT^* (p = 1.9 x 10^-8^). **** = p < 0.0001 using a Mann-Whitney U test. (**E, F**) Confocal live-imaging of *neur-H2B-RFP* (magenta) and *Jupiter-GFP* (green) in a *neur>UAS-mK2-meru^WT^* (**E**) and *neur>UAS-mK2-meru^RxL^* background (**F**). (**G**) Graphs showing the spindle orientation of each genotype relative to the dorsal midline (**G**) and epithelial plane (**G’**). p values indicated for Kolmogorov-Smirnov tests. Scale bars = 10µm. Number of SOPs imaged indicated on the panels from xx animals. Number of SOPs imaged indicated on the panels from 4 animals in panel (**D**) or 3 wild-type or 4 *meru^1^* animals, respectively in panels (**D, F-F’**).

Finally, we analysed the impact of loss of CycA binding on Meru-dependent spindle alignment. mK2-Meru^WT^ expression was sufficient to rescue A-P spindle alignment in the *meru^1^* mutant, however mK2-Meru^RxL^ expression did not rescue A-P spindle angles (Figure 7E-G), though A-B spindle alignment was not significantly affected (Figure 7G’). Thus, CycA docking on Meru is required for posterior Mud recruitment and consequently SOP mitotic spindle alignment.

## Discussion

To build and maintain tissues of the appropriate architecture, cells must read external cues from the extracellular matrix and other cells to align their polarity along (PCP) and across (A-B polarity) the tissue plane (Buckley & St Johnston, 2022; Butler & Wallingford, 2017). A key aspect of tissue organisation is the orientation of the mitotic spindle, which is essential not only to determine the position of the daughter cells after division, but also for cell fate determination through asymmetric cell division (Bergstralh et al., 2017; di Pietro et al., 2016; Lechler & Mapelli, 2021). Although the core PCP and A-B polarity pathways are well conserved, cells adopt radically different polarised organisations and cell division orientations to give rise to the diversity of adult tissues, from the epidermis and intestine to the nervous system (Buckley & St Johnston, 2022; Butler & Wallingford, 2017). Understanding how cells use these conserved polarity complexes as landmarks to generate cell type-specific polarity and spindle orientation remains a key challenge in the field, for which few mechanisms and molecular players have been identified. One of the best understood tissue-specific polarity adaptors is Inscutable (Insc), which was first identified in *Drosophila* neuroblasts (NBs or neural stem cells) (Kraut et al., 1996; Kraut & Campos-Ortega, 1996). In NBs, Insc connects the apical polarity protein Baz with the spindle tethering machinery via Pins, which promotes spindle orientation perpendicular to the tissue plane, allowing the differentiating NB progeny, the ganglion mother cells, to delaminate (Kraut et al., 1996; Schober et al., 1999; Wodarz et al., 1999; Yu et al., 2000).

We had previously shown that Meru, which is transcriptionally activated in SOPs as part of their differentiation programme (Banerjee et al., 2017; Buffin & Gho, 2010; Reeves & Posakony, 2005), links PCP with A-B polarity by promoting Baz planar polarisation (Banerjee et al., 2017). Here, we show that Meru also couples cell polarity with spindle orientation by assembling a ternary complex with the PCP protein Dsh and the spindle tethering factor Mud (Figure 2D), promoting Mud localisation to the posterior cortex to orient the SOP spindle along the A-P axis (Figure 3). In agreement with this model, Mud is depleted from the posterior cortex in *meru* mutants (Figure 3A-C) and the spindle is no longer oriented along the A-P axis (Figure 3D-F’), as in *fz* and *dsh* mutants (Bellaïche, Gho, et al., 2001; Gho & Schweisguth, 1998; Ségalen et al., 2010). Unexpectedly, despite the presence of Meru specifically at the posterior cortex, cortical Mud levels were reduced both anteriorly and posteriorly in *meru* mutant SOPs (Figure 3C). This could indicate that Meru is also required for Mud stability and in its absence, the overall Mud pool is depleted. This is difficult to verify as low Mud expression precludes reliable measurement of its cytosolic levels. However, in our rescue experiments, Meru^WT^ overexpression in a *meru* mutant background causes Mud to be deposited equally at the anterior and posterior poles (Figure 7B’, D), in contrast to wild type SOPs, where we detect more Mud anteriorly than posteriorly (Figure 3C). Together with the fact that Meru^RxL^ expression causes loss of Mud specifically at the posterior pole, this strongly argues in favour of our model that Meru is necessary for posterior Mud recruitment.

Could RASSF proteins such as Meru be implicated in spindle orientation in other contexts? Since the Meru mammalian orthologs RASSF9 and RASSF10 associate with Dvl proteins (Hauri et al., 2013), it would be interesting to test their involvement in oriented cell divisions in contexts in which Dvl1-3 have been implicated, such as HeLa cells (Kikuchi et al., 2010; Yang et al., 2014) and zebrafish gastrulation (Ségalen et al., 2010). Furthermore, similar to *Drosophila* NBs, mouse Inscutable (mInsc) participates in orientating the spindle perpendicular to the tissue plane to promote differentiation of embryonic epidermal progenitors (Lechler & Fuchs, 2005; Poulson & Lechler, 2010; Williams et al., 2014). As RASSF9 is highly expressed in epidermal keratinocytes and *RASSF9* mutant animals show increased proliferation and aberrant differentiation in the epidermis (Lee et al., 2011), it is exciting to speculate that RASSF9 and mInsc could both regulate spindle orientation in this tissue. In *Drosophila*, the Meru paralog RASSF8 is expressed in many epithelial tissues (Langton et al., 2009), therefore it would also be interesting to investigate its role and that of its mammalian orthologs RASSF7 and RASSF8 in symmetric cell division orientation.

Besides positioning, a second important aspect of spindle orientation is temporally linking progression through the cell cycle with spindle tethering (Bergstralh et al., 2017; di Pietro et al., 2016; Lechler & Mapelli, 2021). Several cell polarity and spindle tethering factors such as Lethal giant larvae, Dvl and NuMA, are regulated through phosphorylation by mitotic kinases, supporting the idea of an intimate relationship between spindle orientation and the cell cycle (di Pietro et al., 2016; Lechler & Mapelli, 2021; Osswald & Morais-de-Sá, 2019). We identified a mechanism whereby Mud cortical localisation is coupled to the cell cycle via Meru phosphorylation, linking cell polarity, the cell cycle and spindle orientation. Darnat *et al*. showed that the Cdk1 partner CycA is recruited to the SOP posterior cortex in a Dsh-dependent manner (Darnat et al., 2022). We find that Meru, but not Dsh, associates with CycA in S2 cells (Figure 4C). We uncovered in Meru a pair of overlapping Cyclin-binding RxL SLiMs that are necessary for Dsh/Meru and Mud/Meru binding (Figure 5C and D), the localisation of Meru and Mud to the posterior cortex (Figure 5F, Figure 7C and D), as well as A-P spindle orientation (Figure 7F and G). We therefore propose a model in which, following its posterior recruitment via Dsh, Meru recruits CycA/Cdk1, which in turn phosphorylates Meru, allowing its stabilisation at the posterior cortex and Mud recruitment (Figure 8). Consistent with this model, Meru^RxL^ no longer supports the assembly of the ternary Dsh/Meru/Mud complex (Figure 7A), and a form of Meru mutant for its five Cdk1 consensus sites (Meru^5S/T-A^) displays much reduced Mud binding (Figure 6A).

**Figure 8.**
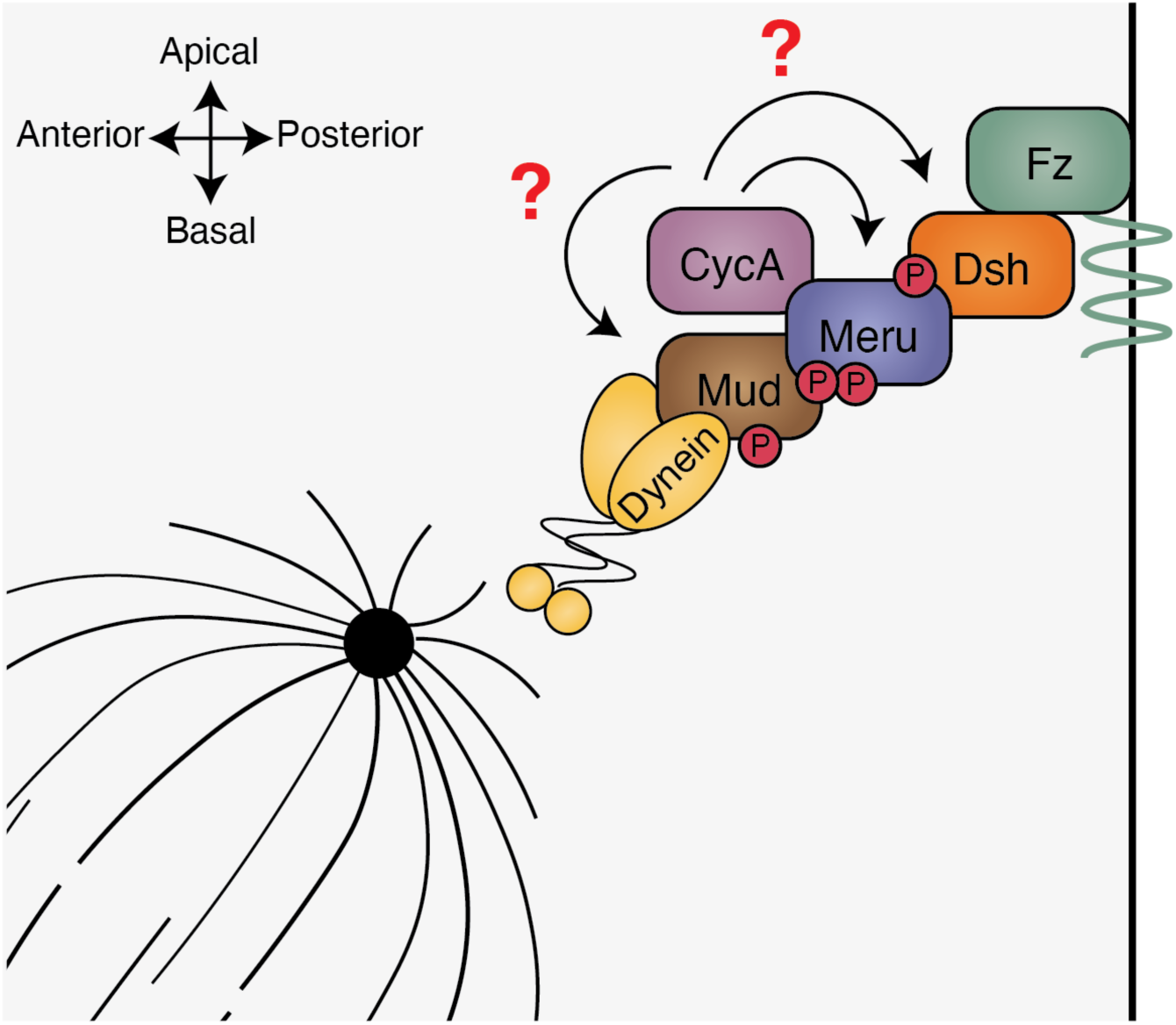
CycA regulates Meru interaction with Mud to tether the mitotic spindle to the posterior cortex of SOPs. Diagram showing our proposed model for Meru function in SOP spindle tethering. Meru, which is recruited to the posterior cortex of SOPs by Dsh, is required to localise Mud. The Meru/Mud interaction is dependent of the docking of CycA to Meru at the 258/9 RxL motif and is regulated by Cdk1 phosphorylation of Meru. Mud and Dsh may also be Meru-dependent CycA/Cdk1 substrates at the SOP posterior cortex.

Does CycA/Meru association promote the phosphorylation of other posterior cortical proteins? Our data indicate that Dsh may be another target, since mutation of the only Cdk1 consensus site in Dsh causes reduced binding to Meru^5S/T-A^ (Figure 6B), suggesting phosphorylation of both partners may be required for optimal binding. It is interesting to note that human Dvl2 phosphorylation on T206 by Plk1 is required for spindle orientation in cultured human cells (Kikuchi et al., 2010), therefore Dsh/Dvl proteins may receive inputs from different cell cycle kinases. Finally, it is possible that Mud is also phosphorylated by Cdk1 in a Meru-dependent manner. Indeed, Cdk1 phosphorylates NuMA on four C-terminal residues to control its cortical turnover in human cells and *C. elegans*, at least in part by inhibiting a NuMA lipid-binding domain (Compton & Luo, 1995; Gehmlich et al., 2004; Hsin-Ling & Ning-Hsing, 1996; Kiyomitsu & Cheeseman, 2013; Kotak et al., 2013; Portegijs et al., 2016; Saredi et al., 1997; Seldin et al., 2013; Zheng et al., 2014).

The cyclin-binding SLiM we identified in the Meru sequence is striking as it is comprised of two overlapping RxL motifs at amino acids 258/259 (Fig EV2). In addition, this site is followed by two optimal Cdk1 consensus phosphorylation sites at amino acid 267 and 275 (Figure 5A). We used this motif to search for proteins containing overlapping RxLs separated by 8 amino acids from a Cdk1 target site ([RK]-[RK]-L-L-x[7]-[ST]-P or RRLL motif). Remarkably, we found that Pins, which recruits Mud to the anterior cortex in SOPs and in many other contexts (Bergstralh et al., 2017; di Pietro et al., 2016; Lechler & Mapelli, 2021), possesses a matching site (Fig EV5A). When we mutated this RRLL motif in Pins, we observed a mislocalisation to the nucleus and delayed localisation to the anterior cortex during mitosis (Fig EV5B and C). As we have shown that Meru binds Mud at the same domain as Pins (Figure 2B), this highlights strong mechanistic parallels between Meru and Pins in terms of Mud recruitment. Pins phosphorylation on S436 by Aurora A increases binding to Dlg (Johnston et al., 2009), suggesting that the Dlg/Pins/NuMA complex is also under tight regulation by the mitotic kinases. Indeed, the Dlg binding partner Guk-holder, which has been implicated in spindle tethering in NBs and S2 cell with induced polarisation (Albertson & Doe, 2003; Garcia et al., 2014; Golub et al., 2017), also has an RRLL motif (Fig EV5A). We found RRLL motifs in other proteins involved in spindle regulation (Aurora borealis (Hutterer et al., 2006), Sister of feo (Vernì et al., 2004)) or cell cycle progression (Double parked (Whittaker et al., 2000)), suggesting it can be used as a strong predictor of regulation by Cdks (Fig EV5A).

In summary, we uncover a mechanism whereby Meru allows coupling of the spindle tethering complex both to the cell polarity machinery and cell cycle progression. We propose that similar regulatory logics underpin the diverse spindle orientation mechanisms employed by other symmetrically and asymmetrically dividing cells.

## Materials and Methods

### Transgenes and fly stocks

*UAS-mK2-meru* transgenic lines were created by phiC31-mediated recombination using stock 9723 from Bloomington (as described in Banerjee *et al*. 2017). The *meru^RxL^* mutant construct was generated under the same protocol but the RxL motif at amino acids 258/9 was mutated to Alanines at sites 258-261. The Pins wild-type and RxL mutant were generated by replacing Meru from the construct above with Pins cDNA and mutating the RxL motif at amino acids 390/1 to Alanines at sites 390-393.

The following fly stocks were used: *neurG4* (Bellaïche, Gho, et al., 2001); *Ecad-GFP* (BL60584); *Dlg-GFP* (VDRC318133); *UAS-Pon-GFP* (Lu et al., 1999); *GFP::mud(62E1), GFP::mud(65B2)* gift from Yohanns Bellaïche (Ségalen et al., 2010); *UAS-cd8-mRFP* (BL27398); *neur-H2B-RFP* (Gomes et al., 2009); *UAS-pins-RNAi* (BL53968); *neurG4, UAS-RFP*, *jupiter::GFP* (recombined by Federica Mangione from BL6825); *meru^1^* (Banerjee et al., 2017); *CycA-GFP*, gift from Michel Gho and Agnes Audibert (Darnat et al., 2022); *UAS-nls-GFP* (BL4716).

### Genotypes

***Fig 1*** (B-C’) w; UAS-mK2-meru/+; neurG4/+ (D) w; Ecad-GFP/UAS-mK2-meru; neurG4/+ (E-E’) w; UAS-mK2-meru/+; neurG4, UAS-Pon-GFP/+ (F-F’’) w; UAS-mK2-meru(WT)/+; GFP::mud(62E1), GFP::mud(65B2), neurG4/+ ***Fig 3*** (A) w; UAS-cd8-mRFP/+ GFP::mud(62E1), GFP::mud(65B2), neurG4/+ (B) w; UAS-cd8-mRFP/+ GFP::mud(62E1), GFP::mud(65B2), neurG4, meru^1^/meru^1^ (D) w;; neur-H2B-RFP, jupiter::GFP (E) w;; neur-H2B-RFP, jupiter::GFP, meru^1^/meru^1^ ***Fig 4*** (A) w; UAS-mK2-meru/+; CycA::GFP, neruG4/+ ***Fig 5*** (F) w;UAS-mK2-meru^RxL^/+; UAS-nls-GFP/neurG4 ***Fig 7*** (B-B’) w; UAS-mK2-meru^WT^/+; GFP::mud(62E1), GFP::mud(65B2), neurG4, meru^1^/meru^1^ (C-C’) w; UAS-mK2-meru^RxL^/+; GFP::mud(62E1), GFP::mud(65B2), neurG4, meru^1^/meru^1^ (E) w; UAS-mK2-meru^WT^/+; neurG4, UAS-RFP, jup::GFP, meru^1^/meru^1^ (F) w; UAS-mK2-meru^RxL^/+; neurG4, UAS-RFP, jup::GFP meru^1^/meru^1^ **Fig EV1** (A-A’) w; Ecad-GFP/UAS-mK2-meru; neurG4/+ (B-B’) w; UAS-mK2-meru(WT)/+; GFP::mud(62E1), GFP::mud(65B2), neurG4/+ (D-D’) w; UAS-mK2-meru/+; neurG4/+ **Fig EV3** (C-C’) w; Ecad-GFP/UAS-mK2-meru^RxL^; neurG4/+ (D-D’) w; UAS-mK2-meru^RxL^/+; neurG4, Dlg-GFP/+ **Fig EV5** (b) w; UAS-mK2-Pins^WT^/+; neurG4/+ (C) w; UAS-mK2-Pins^RxL^/+; neurG4/+ **Movie EV1** w;; neur-H2B-RFP, jupiter::GFP **Movie EV2** w;; neur-H2B-RFP, jupiter::GFP, meru^1^/meru^1^

### Confocal live-imaging

*Drosophila* pupae were attached to a Superfrost microscope slide (Thermo Scientific) using doubled sided tape so that the A-P axis was positioned along the width of the slide. Once adhered, the pupal case from the head to the start of the abdomen was peeled off. Two stacks of four 22 x 22mm No. 1.5 Cover Glass (VWR) were glued together and placed on either side of the pupae. A lightly oiled 24 x 50 mm No. 1.5 Cover Glass (VWR) was placed atop the two cover slip stacks so that the oil side rested gently on the exposed notum. The sample was then inverted to image on an invert Nikon Spinning Disk (60x or 100x oil objectives) confocal microscope.

### Adult notum and wing imaging

Female adults, raised at 25°C, were collected based on genotype and were stored at −20°C or in 70% EtOH at 4°C for notum and wing imaging, respectively. Adult nota were mounted in 22 0.8% low melting agarose and imaged on an MZ16 stereomicroscope (Leica) with a QImaging MicroPublisher 6 Color Camera. Adult wings were mounted onto Superfrost microscope slides (Thermo Scientific) in Euparal covered in 22 x 22mm No. 1.5 Cover Glass (VWR) for imaging. Wings were imaged with an Axio Scan.Z1 (Zeiss) slide scanner with a 2.5x and 10x objective.

### Colocalisation analysis, cortical Mud levels and spindle orientation in time-lapse movies

All analyses were carried out using ImageJ. The degree of correlation between Meru and Mud/CycA localisation at the posterior cortex was measured by drawing a 2 μM thick line over the Meru cortical crescent. The line was applied to the single brightest slice (0.6 µm thick) in the stack used in the corresponding figure to the Meru and Mud/CycA channels, and the normalised grey value was plotted versus the distance along the line (in μm). The correlation between the Meru and Mud/CycA gray value levels was calculated by the Pearson’s correlation coefficient (r) in which 1 represents perfect correlation, −1 represents anti-correlation and 0 depicts no correlation.

Levels of cortical Mud were quantified by measuring the sum of the integrated density of a cortex-specific ROI corrected for background fluorescence and ROI area. A mask of the membrane marker (CD8-mRFP) was used to outline the cell membrane and placed over the GFP channel to isolate the signal to the dividing cell. As the mitotic spindle in SOPs has a z-tilt (David et al., 2005), the anterior and posterior crescents were analysed separately. The three brightest slices (0.6 µM z-slices) for GFP::Mud signal were max-projected and isolated by drawing around the ROI, ensuring that the signal coming from the centrosome was excluded. The cytoplasmic signal (background fluorescence) was measured using a 10×10 pixel square for each cortical max-projection. The sum of the anterior and posterior cortical Mud signals, respectively, were corrected for background fluorescence and area of the ROI by the following formula:

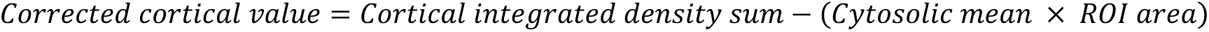

The anterior-posterior (A-P) ratio was found with the resulting values:

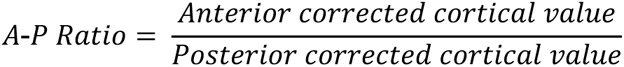

Movies obtained by confocal microscopy were imported into the Imaris Microscopy Image Analysis Software (Oxford Instruments) to quantify the x- and z-axis of every centrosome pair in dividing cells of interest. All measurements were made by manually positioning a point at the center of either centrosome, creating a line through the spindle. The x-, y-, and z-coordinates of each point were used to quantify the angle of division relative to a reference position. The first measurement pair was used to define the dorsal midline, identified by visual cues such as anisotropic cell shape, followed by measurement pairs at all dividing cells of interest. The spindle angles in x-axis (A-P axis) were calculated as degrees away from the midline. The reference slope (the midline angle) was calculated by the following equation:

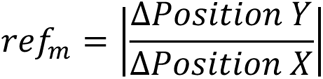

To quantify the x-axis angle of each centrosome pair, the slope was found with the same equation above (called centrosome_m_) and calculated as follows:

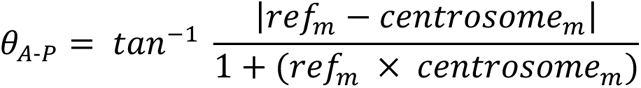

The z-axis angle (A-B axis) was calculated as degrees away from the plane. The reference plane angle was set to 0. and the absolute z-axis angle of each centrosome pair was calculated with the following equation:

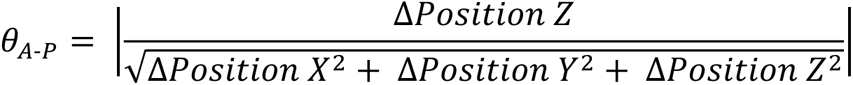

### Statistical analysis

Mann-Whitney U tests were performed to test if mean values of cortical Mud levels were significantly different. The Kolmogorov-Smirnov (K-S) test was used to test if measurements from SOP spindle alignment samples distributions were significantly different. To test the colocalisation of two fluorescently-tagged proteins in a time-lapse movie, the Pearson’s correlation test was performed.

### *Drosophila* cell culture and expression constructs

*Drosophila* S2 cells were transfected with Effectene transfection reagent (Qiagen) and grown in *Drosophila* Schneider’s medium (Life Technologies) containing 10% FBS (Sigma), 50μg/ml penicillin and 50 μg/ml streptomycin. Expression plasmids were generated using Gateway® technology (Life Technologies). Open reading frames (ORF) were PCR amplified from existing plasmids or cDNA (DGRC, https://dgrc.cgb.indiana.edu/vectors/) and cloned into entry vectors (pENTR^TM^/D-TOPO). Vectors from the *Drosophila* Gateway Vector Collection (http://www.ciwemb.edu/labs/murphy/Gateway%20vectors.html) were used as Destination vectors. Meru, Mud and Baz were tagged N-terminally and Dsh, Cdk1 and all the Cyclins were tagged C-terminally. Point mutations were generated using the Quikchange Multi Site-Directed Mutagenesis Kit (Agilent). All expression plasmids were sequence-verified.

### dsRNA production and treatment

dsRNAs were synthesized using the Megascript T7 kit (Thermo Fisher Scientific). DNA templates for dsRNA synthesis were PCR-amplified from genomic DNA using primers that contained the 5’ T7 RNA polymerase-binding site sequence. dsRNAi primers were designed using the DKFZ RNAi design tool (http://www.dkfz.de/signaling2/e-rnai/). RNAi experiments were carried out in 6 well plates using S2 cells. 1-2×10^6^ cells were plated per well and cells left to settle. After 3h, the medium was removed and replaced with 1 mL of serum free Schneider’s medium containing dsRNAs (20 μg). Cells were soaked for 30 mins and then 2 mL of full Schneider’s medium was added. Transfections were carried out 1h after dsRNA treatment.

### Immunoprecipitation and immunoblot analysis

For immunoprecipitation assays, cells were lysed in lysis buffer (150 mM NaCl, 50 mM HEPES pH 7.5, 0.5% (v/v) Triton X-100 supplemented) supplemented with phosphatase inhibitor cocktails 2 and 3 (Sigma) and protease inhibitor cocktail (Roche). Cell extracts were cleared at 13,200 rpm for 20 mins at 4°C. GFP and Myc-tagged proteins were purified using agarose beads (Chromotek) according to the manufacturers protocol. Detection of purified proteins and associated complexes was performed by immunoblot analysis using chemiluminescence (Pierce). Western Blots were probed with mouse anti-FLAG (M2, Sigma), rabbit anti-FLAG (Sigma), mouse anti-Myc (9E10, Santa Cruz Biotechnology), rabbit anti-HA (C29F4, NEB), mouse anti-Tubulin (E7, Developmental Studies Hybridoma Bank, DSHB) and rat anti-GFP (Chromotek).

## Acknowledgements

We thank Y. Bellaïche, M. Gho, F. Schweisguth, F. Mangione and the Bloomington *Drosophila* Stock Center for fly stocks and Y. Bellaïche for plasmids. We are grateful to M. Renshaw and members of the Crick Advanced Light Microscopy Facility for advice on confocal imaging and image analysis; J. Kurth, S. Murray, H. Shaw, S. Ruiz-Herrera from the Crick Fly Facility for embryo injections and fly stock maintenance; and Tapon lab members for advice. We thank A. Moraiti, M. Holder, F. Mangione, N. Goehring and A. Audibert for critical reading of the manuscript. This work was supported by the Francis Crick Institute, which receives its core funding from Cancer Research UK (CC2138), the UK Medical Research Council (CC2138) and the Wellcome Trust (CC2138) and a Wellcome Trust Investigator award (107885/Z/15/Z) to N.T. For the purpose of Open Access, the authors have applied a CC BY public copyright licence to any Author Accepted Manuscript version arising from this submission.

## Author Contributions

Conceptualization, all authors; funding acquisition, N.T.; experiments and data analysis, M.M., B.L.A, J.J.B.D.; writing – original draft, N.T., M.M., B.L.A; writing – review & editing, all authors.

## Expanded View

**Figure EV1.**
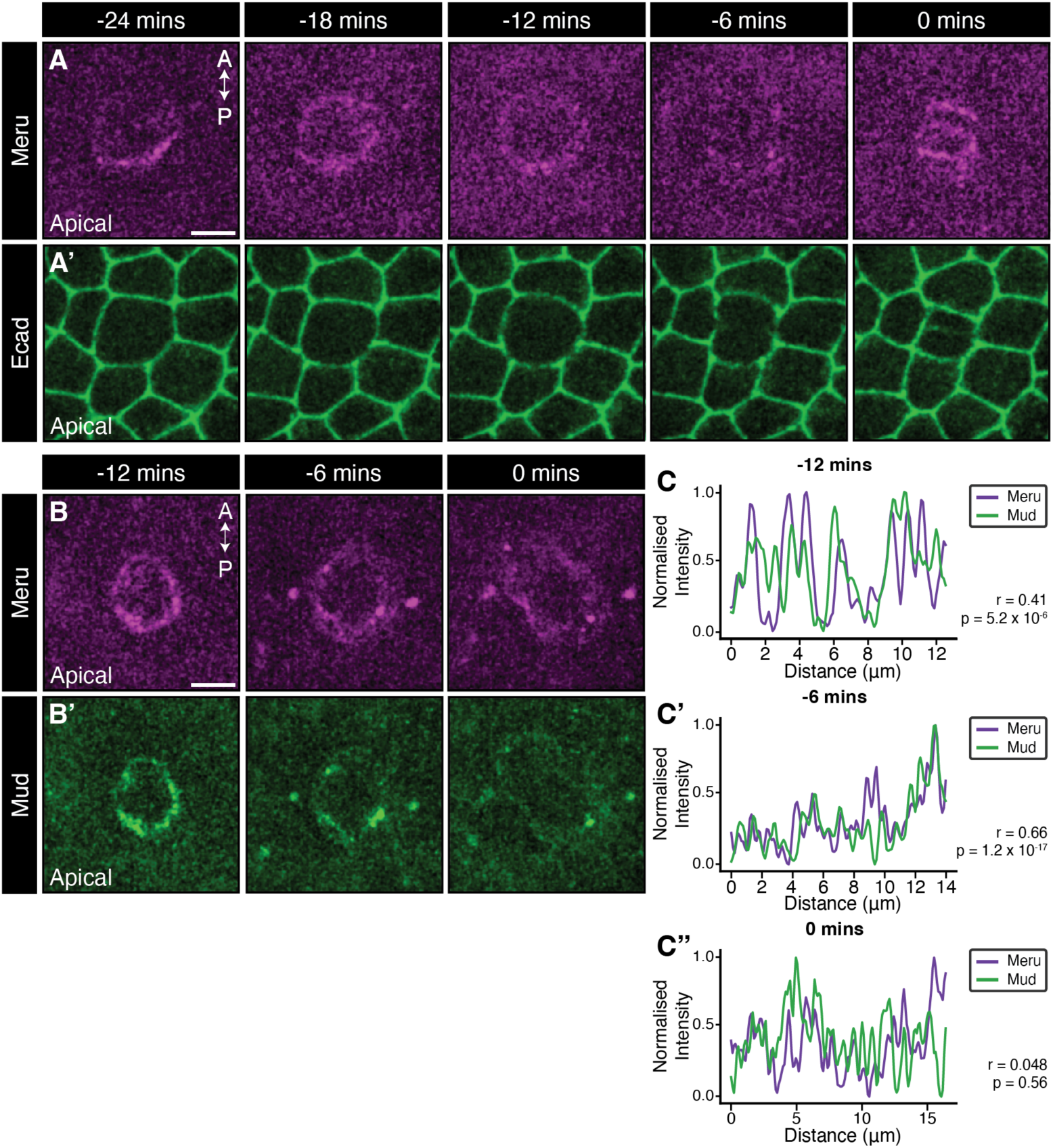
Posterior-apical localisation of Meru occurs prior to SOP mitosis and persists throughout division. **(A, B)** Maximum intensity projections of pupal nota live-imaged on a confocal microscope at 16 h APF, 0 mins marking the first frame indicating cytokinesis, of *neurG4>UAS-mK2-meru* (magenta; panels **A** and **B**), co-expressed with *Ecad-GFP* and *Mud-GFP* (green; panels **A’** and **B’**, respectively). (**C-C’’**) Single brightest slice in panels **B** and **B’** plotted as line graphs by measuring the grey value across the posterior domain as defined by Meru localisation normalised to highest value in each channel. High Pearson’s coefficient (*r*) indicates positive correlation. p-values calculated using a two-tailed test. Scale bars = 5 µm.

**Figure EV2.**
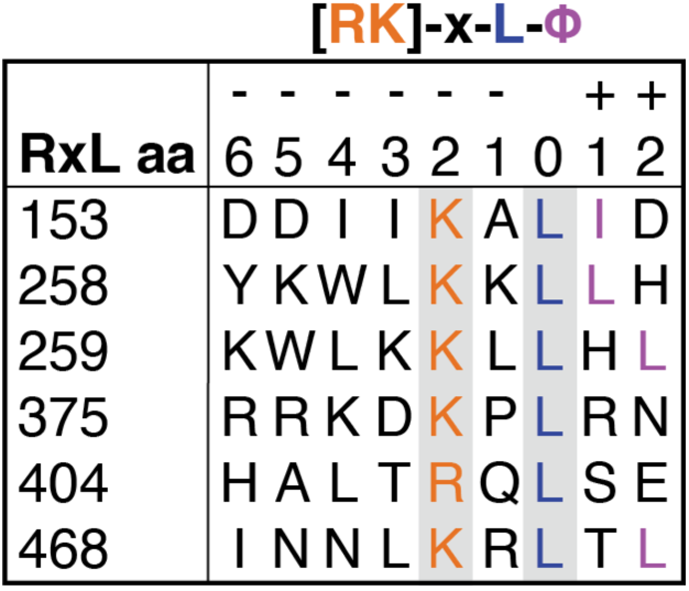
The RxL motifs of Meru. The amino acid (aa) positions of the six RxL motifs identified in the Meru sequence are indicated on the left. On the right are the sequence alignment of each RxL motif. Orange indicates the R/K and the L is labelled in blue at position 0. The hydrophobic aa is marked in purple and the grey boxes indicate the two aa that were mutated to Alanine in RxL mutants.

**Fig EV3.**
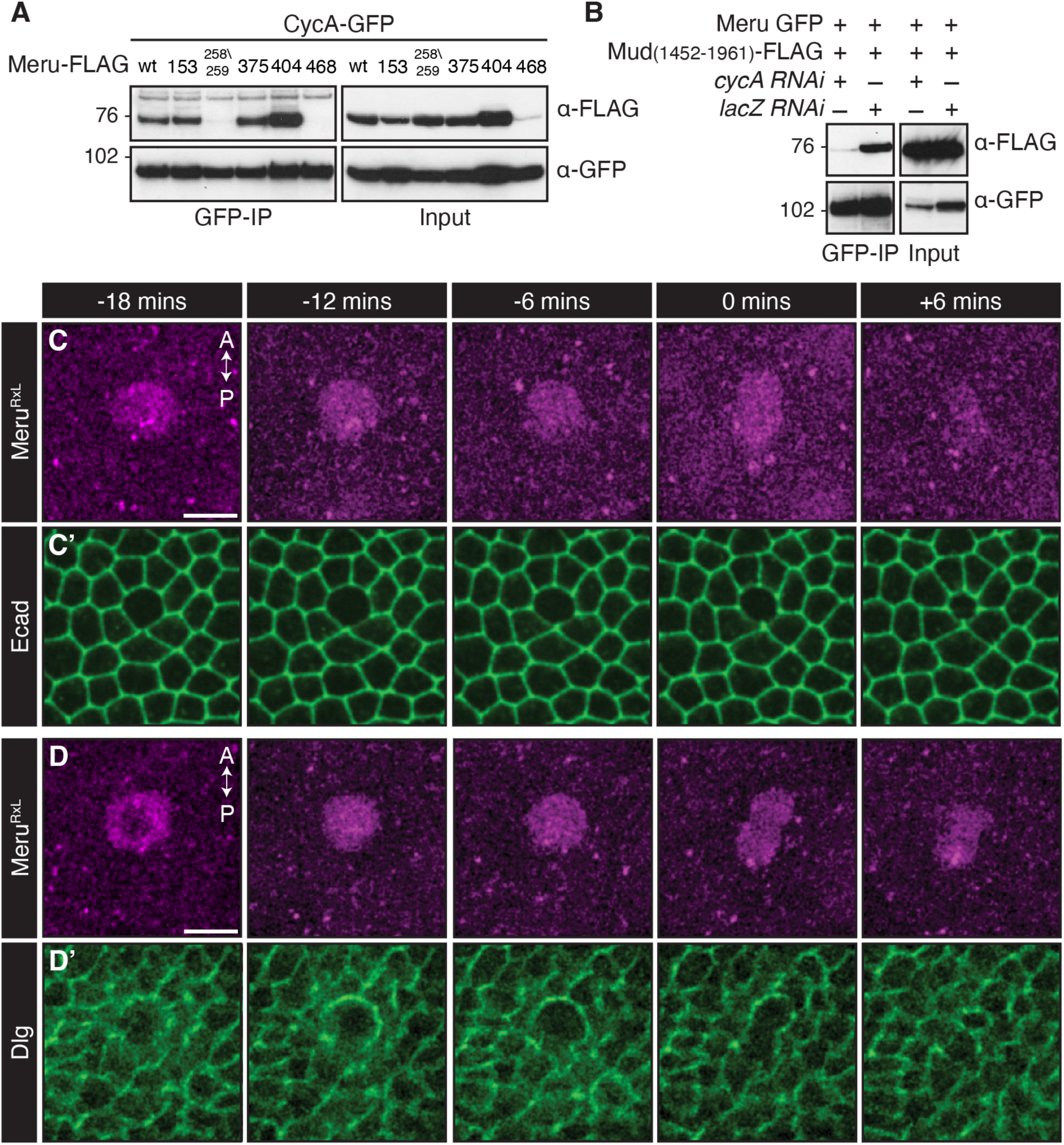
258/9 RxL motif mutant Meru no longer binds CycA and is mislocalised *in vivo*. (**A, B**) Western blots of co-IP experiment using cell lysates from transfected S2 cells, immunoprecipitated and probed using the indicated antibodies. (**A**) CycA immunoprecipitates with each Meru RxL motif mutant, except when Meru 258/9 is mutated to Alanines. (**B**) CycA depletion by RNAi reduced Meru/Mud association. (**C, D**) Maximum intensity projections of pupal nota live-imaging at 16 h APF, 0 mins marking the first frame indicating cytokinesis, of *neurG4>UAS-mK2-meru^RxL^* (magenta; panels **C** and **D**), co-expressed with apical *Ecad-GFP* and basolateral *Dlg-GFP* (green; panels **C’** and **D’**, respectively). Scale bars = 10 µm.

**Figure EV4.**
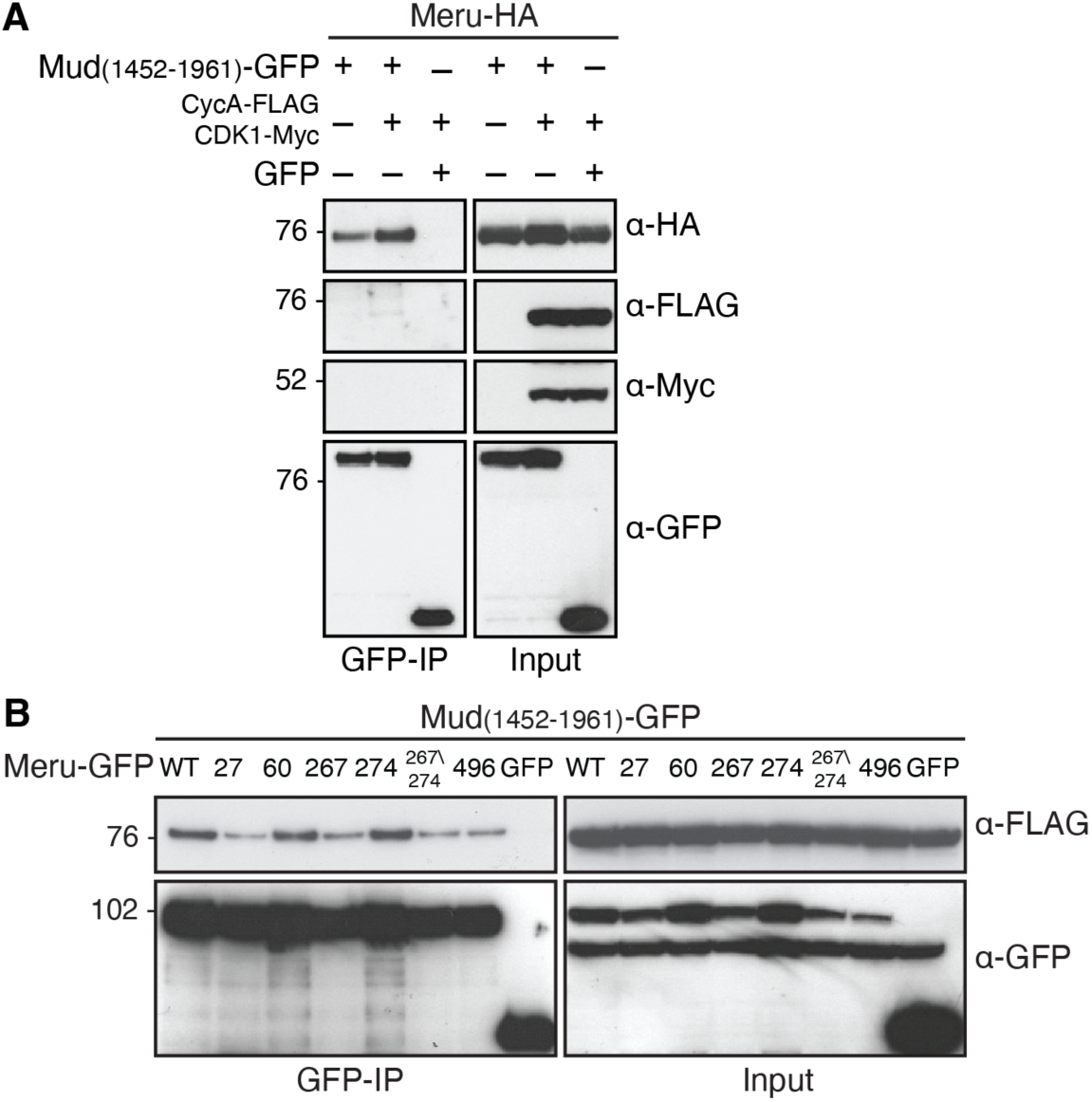
Overexpression of Cdk1 and CycA promotes the Meru/Mud interaction. (**A, B**) Western blots of co-IP experiment using cell lysates from transfected S2 cells, immunoprecipitated and probed using the indicated antibodies. CycA immunoprecipitates with C-terminal Mud in the presence of Meru (**A**) and no single Meru S/T-P phosphosite is essential to co-IP with C-terminal Mud **(B)**.

**Figure EV5.**
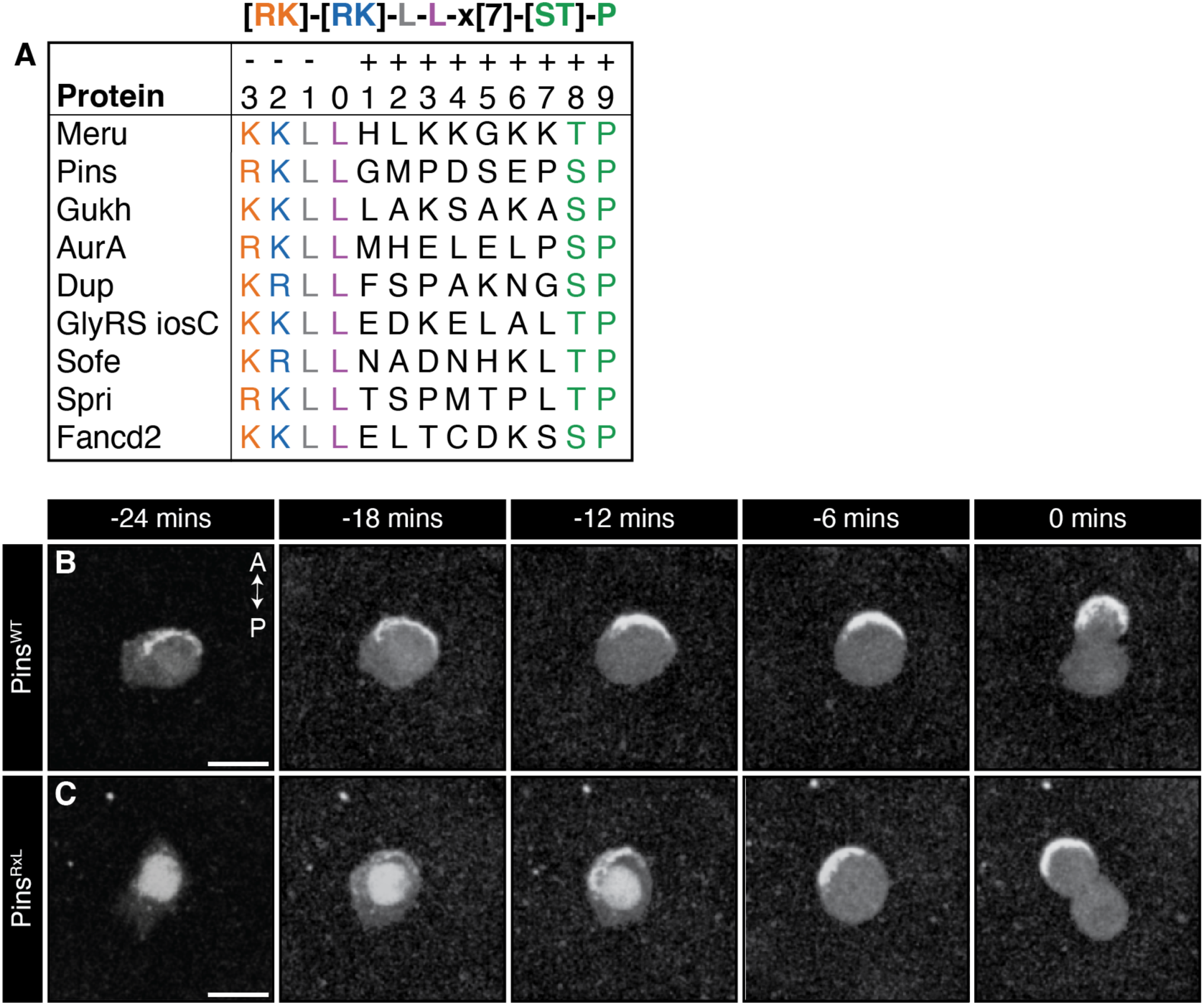
Loss of the Pins RxL motif results in delayed cortical localisation. **(A)** BLAST search results for novel [RK]-[RK]-L-L-x[7]-[ST]-P (RRLL) motifs resulted in nine proteins of interest, including Meru, Pins, Gukh and AurA, all of which are involved in spindle orientation. (**B, C**) Confocal live-imaging of pupal nota at 16 h APF, 0 mins marking first frame in cytokinesis, shows expression of *UAS-mK2-Pins^WT^* (**B**) and *UAS-mK2-Pins^RxL^* (**C**) driven by *neruG4* during SOP mitosis. Scale bars = 10 µm.

**Movie EV1**

**Title:** Wild-type SOPs divide along the A-P axis

**Description:** Confocal live-imaging of an SOP division in the pupal notum at 16 h APF. SOPs are marked by *neur-H2B-RFP* (magenta) and the spindle is marked by *jup::GFP* (green). The spindle aligns along the A-P axis and the daughter cells are segregated anteriorly and posteriorly (top and bottom of frame, respectively). Scale bar = 10 μm

**Movie EV2**

**Title:** Loss of *meru* leads to spindle misalignment during SOP division

**Description:** Confocal live-imaging of an SOP division in the *meru^1^* mutant background in the pupal notum at 16 h APF. SOPs are marked by *neur-H2B-RFP* (magenta) and the spindle is marked by *jup::GFP* (green). In this example, the spindle aligns perpendicular to the A-P axis (top to bottom of frame). Scale bar = 10 μm.

